# Emerging fish pathogens *Lactococcus petauri* and *L. garvieae* in Nile tilapia (*Oreochromis niloticus*) farmed in Brazil

**DOI:** 10.1101/2022.08.19.504548

**Authors:** Renata Catão Egger, Júlio César Câmara Rosa, Santiago Benites de Pádua, Fernanda de Oliveira Barbosa, Mariana Taíse Zerbini, Guilherme Campos Tavares, Henrique César Pereira Figueiredo

**Author notes:** Corresponding author: Henrique César Pereira Figueiredo, Department of Preventive Veterinary Medicine, School of Veterinary Medicine, Federal University of Minas Gerais, Belo Horizonte, Brazil. Zipcode: 31270-901.

## Abstract

Lactococcosis in fish has been associated with *Lactococcus garvieae* and the recently described *L. petauri*. However, the relevance of these emerging fish pathogens to Nile tilapia still requires thorough understanding. This study investigated lactococcosis outbreaks in Nile tilapia on Brazilian farms and characterized the isolates through molecular identification of the bacterial species, multilocus sequence typing (MLST) analysis, virulence to Nile tilapia, and antimicrobial susceptibility. Lactococcosis outbreaks were monitored from 2019 to 2022 throughout Brazil. The outbreaks occurred mainly during warmer months, and co-infections were observed in four farms, whereas concurrent bacterial infections were identified in all farms. Since the sequence of the 16S rRNA was not capable of differentiating between *L. petauri* and *L. garvieae*, *Lactococcus* spp. isolates were identified at the species level using the *gyrB* gene sequence. In total, 30 isolates were classified as *L. petauri* and two as *L. garvieae*. All *L. petauri* isolates were grouped in ST24, except for one isolate which belonged to the newly described ST47. A new ST was also described for the *L. garvieae* isolates identified, ST46. Furthermore, *L. petauri* ST24 and ST47 were characterized as singletons, whereas *L. garvieae* ST46 was grouped with ST16 and ST17 and formed CC17. For the challenge trial, an *L. petauri* ST24 isolate was chosen considering that this MLST lineage was the most frequently observed. *L. petauri* was reisolated from challenged Nile tilapia, confirming the pathogenicity of this bacterium to Nile tilapia. The infection in the fish progressed very rapidly, and within 48 h post-challenge clinical signs and the first mortalities were observed. The estimated LD50 was 5.74 × 10^3^ CFU 15 days post-challenge. Provisional epidemiological cutoff values were determined for *L. petauri* for six antimicrobial agents from different drug classes. All isolates were characterized as wild type (WT) for neomycin and oxytetracycline, whereas 96.67 % of the isolates were characterized as WT for amoxicillin, erythromycin, and florfenicol, and 83.33 % were WT for norfloxacin. Of the 14 outbreaks analyzed, 12 were caused by *L. petauri* and two by *L. garvieae*. The *gyrB* gene sequence was used to differentiate *L. petauri* from *L. garvieae* and allowed for the correct identification of these pathogens. Two MSLT lineages of *L. petauri* were identified and ST24 was observed in different regions of the country, illustrating a rapid expansion of this bacterial lineage.

**Highlights of the manuscript:**

- *Lactococcus petauri* is pathogenic to Nile tilapia.
- The MLST lineage most observed was *L. petauri* ST24, indicating its adaption to infect Nile tilapia.
- The analysis of the *gyrB* gene sequence allowed for the correct identification of *L. petauri* and *L. garvieae*.

## 1 Introduction

Lactococcosis is an emergent bacterial disease that causes hemorrhagic septicemia in fish. The etiological agent which has been associated with piscine lactococcosis is *Lactococcus garvieae* (Bwalya et al., 2020b; Nishiki et al., 2016; Ortega et al., 2020; Rao et al., 2021). This bacterium infects a wide range of animal hosts (Evans et al., 2006; Kawanishi et al., 2006; Morick et al., 2022; Pot et al., 1996; Teixeira et al., 1996; Tejedor et al., 2011; Thiry et al., 2021) and is also considered as a potential foodborne zoonotic agent with clinical significance for humans (Chan et al., 2011; Russo et al., 2012; Wang et al., 2007).

More recently, *Lactococcus petauri* was described as a novel species following a *L. garvieae* subgroup A genomic reassignment (Goodman et al., 2017). Importantly, *L. petauri* has been associated with lactococcosis in fish, as some isolates obtained from diseased fish and previously identified as *L. garvieae* have been reclassified as *L. petauri* (Altinok et al., 2022; Kotzamanidis et al., 2020; Shahin et al., 2022). *Lactococcus garvieae* and *L. petauri* seem to cause similar clinical signs in fish and both bacterial pathogens are responsible for significant economic losses due to high fish mortality, reduced growth, and treatment costs during outbreaks on fish farms.

Lactococcosis has been reported in different aquaculture species, such as Nile tilapia (*Oreochromis niloticus*), rainbow trout (*Oncorhynchus mykiss*), yellowtail (*Seriola quinqueradiata*), amberjack (*Seriola dumerili*), and cobia (*Rachycentron canadum*) (Bwalya et al., 2020b; Karami et al., 2019; Nishiki et al., 2011; Rao et al., 2021). In Brazil, *L. garvieae* infection has been reported in Nile tilapia and in catfish (*Pseudoplatystoma* sp.) (Evans et al., 2009; Fukushima et al., 2017; Sebastião et al., 2015; Tavares et al., 2018). Infections of *L. garvieae* in Nile tilapia have been more frequently diagnosed in the last years, and this pathogen has consequently become more relevant (Anshary et al., 2014; Bwalya et al., 2020b; Evans et al., 2009; Osman et al., 2017; Tsai et al., 2012). However, most cases were reported before *L. petauri* was described; thus, the relevance of *L. petauri* as a bacterial pathogen for this fish species should be further assessed. The increase and spread of Nile tilapia farming worldwide (FAO, 2022), together with the intensive farming of this fish create a dynamic scenario of pathogens associated with Nile tilapia, which requires close monitoring.

The clinical signs of *L. garvieae* infection in Nile tilapia include erratic swimming, decreased uptake of food, lethargy, dark skin pigmentation, exophthalmia, corneal opacity, intraocular hemorrhage, and body wound/ skin ulcer (Bwalya et al., 2020a, 2020b; Osman et al., 2017). The reported outbreaks occurred during warmer months and mortalities of up to 18 % have been reported in fish > 200 g (Bwalya et al., 2020b). Despite the increasing importance of *L. garvieae* in Nile tilapia production, information on the infection by this emerging pathogen is scarce. Similarly, when *L. petauri* is considered, knowledge on this subject requires further development.

Therefore, in this study, we report outbreaks of *L. petauri* and *L. garvieae* in Nile tilapia on Brazilian farms and characterize the isolates obtained. This study aimed to characterize lactococcosis outbreaks through the identification of the bacterial species involved, and analysis of *Lactococcus* spp. genotyping, virulence, and antimicrobial susceptibility.

## 2 Material and methods

### 2.1 Outbreaks presentation and fish sampling

The first outbreak occurred in a Nile tilapia farm in October 2020 in the state of Mato Grosso, Brazil, resulting in mortalities above 15% in several floating cages. The producer reported erratic swimming near the water surface, exophthalmia, dark skin pigmentation, and eroded gills. Sixty-six diseased fish were sampled (average weight of 715.11 g ± 339.88) and transported on ice to the laboratory for analysis. Complete postmortem examinations were performed on the fish and samples were collected for bacteriological analysis.

After this outbreak, other cases of lactococcosis on Nile tilapia in Brazilian fish farms were monitored and investigated. Outbreaks occurred on Nile tilapia farms located in different regions of the country (Figure 1) from 2020 to 2022. Fish exhibited signs of lethargy, erratic swimming, exophthalmia, corneal opacity, dark skin pigmentation, necrotic and/or pale gills, eroded fins and coelomic distention (Figure 2). Internally, the fish exhibited liver, kidney, spleen pallor, and hemorrhages were observed in the liver and intestine. In addition, an isolate from an outbreak that occurred in 2019 and was previously identified was included in this investigation. In total, 12 fish farms were sampled, of which two had history of vaccination against *Streptococcus agalactiae* serotype Ib (farms 2 and 12), three vaccinated against *S. agalactiae* serotypes Ib and III (farms 4, 7 and 8), one vaccinated against *S. agalactiae* serotypes Ia, Ib and III (farm 5), one vaccinated against *S. agalactiae* serotypes Ib and III + *Streptococcus dysgalactiae*, and other four farms did not use any vaccines (farms 1, 3, 6, 9, 10 and 11).

**Figure 1.**
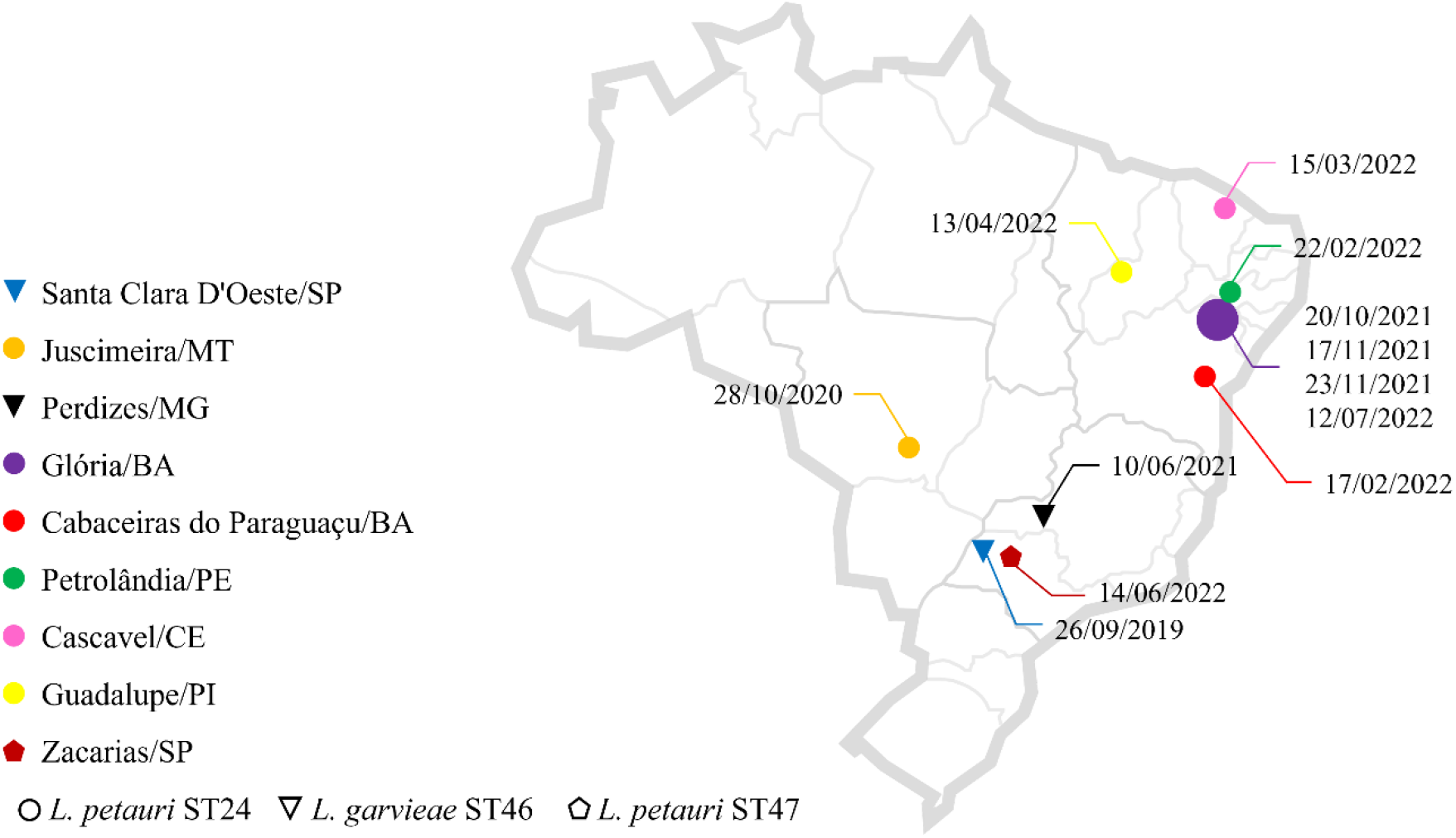
Map of the distribution of *L. petauri* and *L. garvieae* sequence types (ST) according to farms geographic location and sampling date throughout Brazil. Symbols sizes are proportional to the number of farms on that location.

**Figure 2.**
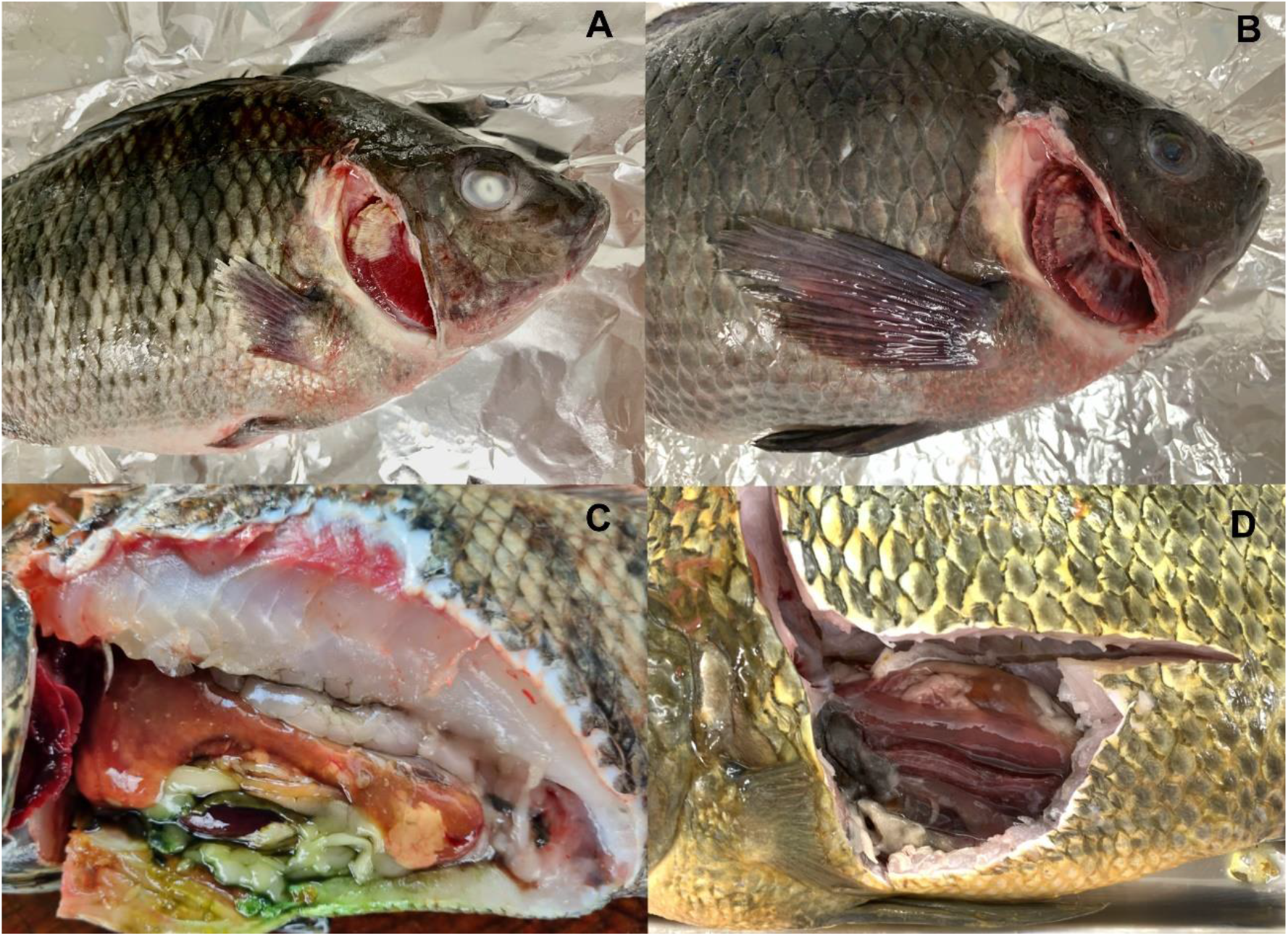
Pictures of Nile tilapia infected with *L. petauri* exhibiting gill necrosis (A and B), exophthalmia (A) corneal opacity (A and B), eroded fin (A), skin petechial hemorrhages (A and B), liver pallor with hemorrhagic areas (C and D), and hemorrhagic intestine (D).

Two *Lactococcus* spp. isolates from each of these outbreaks were selected and characterized according to species identification, multilocus sequence typing (MLST), and antimicrobial susceptibility. These isolates are presented in Table 1.

**Table 1.**
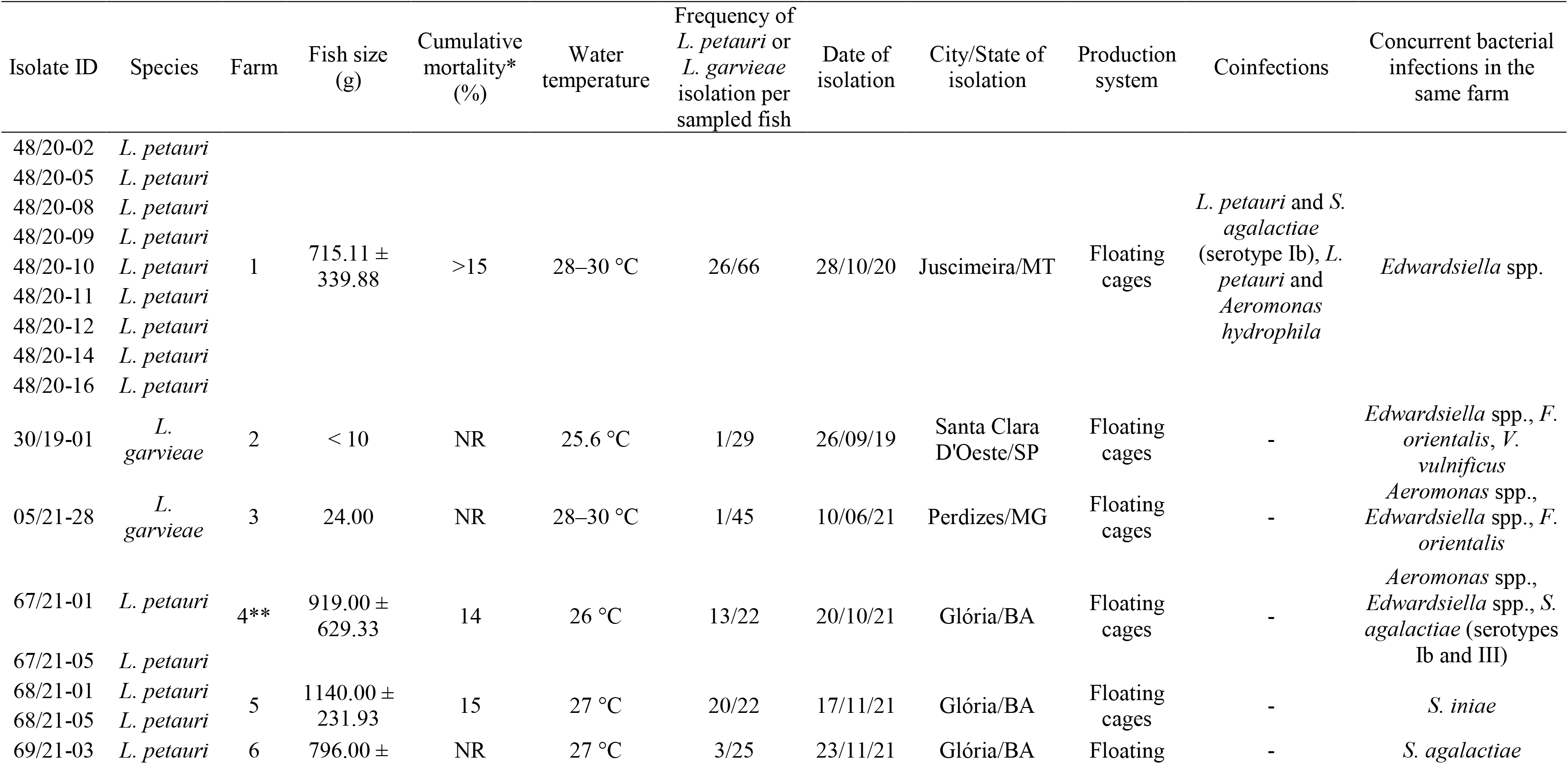
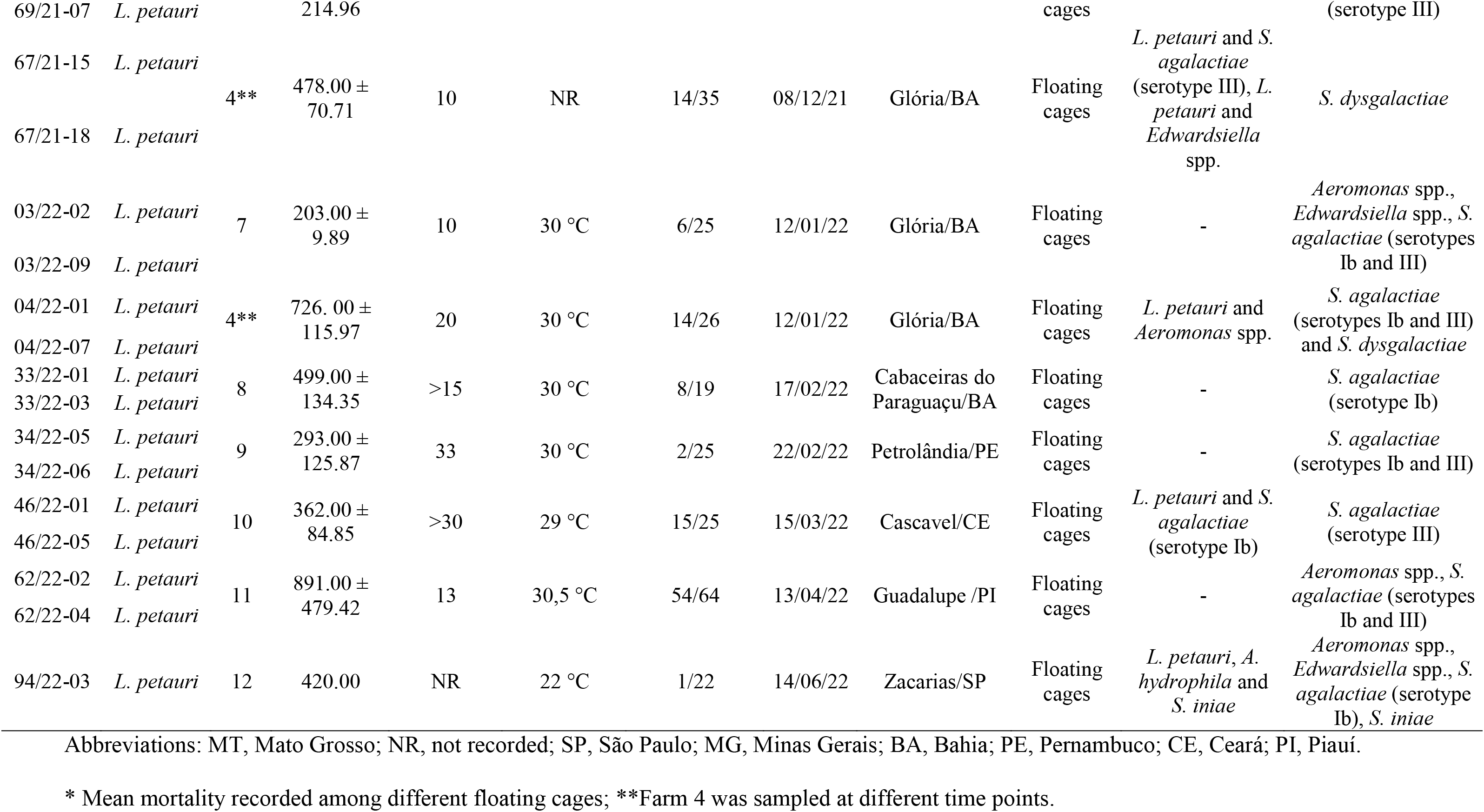
*Lactococcus petauri* and *L. garvieae* outbreaks in Brazil between in 2020 and 2022

### 2.2 Bacterial isolation and identification

During outbreaks investigation, swabs of the brain and kidney were aseptically obtained from the fish, streaked onto Tryptic Soy Agar (TSA) (HiMedia, Mumbai, India) supplemented with 5 % sheep blood and swabs of spleen and kidney were streaked onto Cysteine Heart Agar (HiMedia) supplemented with 1 % bovine hemoglobin and incubated at 28 °C for 2–7 days to isolate bacterial pathogens. Bacterial species were identified using matrix-assisted laser desorption-ionization time-of-flight (MALDI-TOF) mass spectrometry with a Bruker Microflex MALDI Biotyper 2.0 (Bruker Daltonics, Billerica, MA, USA), as previously described (Assis et al., 2017). After the identification of the bacterial species, isolates identified as *Lactococcus* spp. were stored at −80 °C in brain heart infusion (BHI) (HiMedia) broth supplemented with 15 % (v/v) glycerol.

#### 2.2.1 Molecular identification of bacterial isolates at species level

For molecular analysis, *Lactococcus* spp. isolates were thawed from freezer stocks and grown on TSA supplemented with 5 % sheep blood at 28 °C for 24 h. Colonies were collected and subjected to total DNA extraction using the Maxwell® 16 Tissue DNA Purification Kit in the Maxwell 16 Research Instrument (Promega, Madison, WI, USA) following the manufacturer’s instructions. Extracted DNA was quantified using a NanoDrop spectrophotometer (Thermo Fisher Scientific, Waltham, MA, USA) and stored at −80 °C until use.

For 16S rRNA gene sequencing, amplifications were performed by PCR with the universal primers C70 (5’-AGA GTT TGA TYM TGG C-3’) and B37 (5’-TAC GGY TAC CTT GTT ACG A-3’) according to the method described by Fox et al. (1995) with slight modifications. The primers used were purchased from Integrated DNA Technologies (IDT, Coralville, IA, USA). Briefly, PCR reactions were performed with a HotStart Taq polymerase kit (Qiagen, Germantown, MD, USA) in a final reaction volume of 38 μL. The reaction mixture consisted of 1× PCR buffer, 0.25 μM of each primer, 0.2 mM dNTPs, 0.5 mM MgCl_2_, 0.0625 U/μL Taq DNA polymerase and 50 ng DNA template. The PCR conditions consisted of an initial denaturation step at 95 °C for 15 min followed by 35 cycles consisting of 95 °C for 1 min, 58 °C for 1 min, and 72 °C for 1 min, with a final extension step at 72 °C for 15 min. The PCR was performed in a Veriti 96-Well Thermal Cycler (Thermo Fisher Scientific), and amplicons were separated in QIAxcell Advanced using the QX DNA Screening Kit (Qiagen).

Subsequently, amplicons were purified using Agencourt AMPure XP (Beckman Coulter, USA), following the manufacturer's instructions, and sequenced using forward and reverse primers. Sequencing reactions were performed using a BigDyeTM Terminator Cycle Sequencing Kit (Applied Biosystems, Foster City, CA, USA) and run on an ABI 3500 Genetic Analyzer (Applied Biosystems). Sequences were then compared to sequences from the NCBI database using the BLASTN algorithm.

Since the 16S rRNA sequence was not sufficiently different to discriminate between *L. petauri* and *L. garvieae*, the sequence of the gene *gyrB* was used for phylogenetic analysis of the isolates (Shahin et al., 2022). Forward and reverse sequencing products of *gyrB* from the MLST analysis (described in the next section) were used to generate consensus sequences (Supplementary Table S1). The available *gyrB* sequences from *L. petauri* and *L. garvieae* isolates whose identities have been previously confirmed at the species level were then downloaded from NCBI and all sequences were aligned using MEGA v.11 software (Tamura et al., 2021). Evolutionary distances were computed using the Maximum Composite Likelihood method (Tamura et al., 2004), and evolutionary history was inferred using the Neighbor-Joining method (Saitou and Nei, 1987). A bootstrap analysis to investigate the stability of the trees was performed on 1,000 replicates (Felsenstein, 1985).

### 2.3 MLST and eBURST analysis

The MLST scheme for *L. garvieae* was performed to analyze the population structure of *L. petauri* and *L. garvieae* isolates. The isolates were sequence typed using seven housekeeping genes: *als* (α-acetolactate synthase), *atpA* (α-subunit of ATP synthase), *tuf* (elongation factor EF-Tu), *gapC* (glyceraldehyde-3-phosphate dehydrogenase), *gyrB* (DNA gyrase β-subunit), *rpoC* (RNA polymerase β’-subunit), and *galP* (galactose permease) (Ferrario et al., 2013). The genes were amplified by PCR as described previously (Ferrario et al., (2013). Briefly, PCR was performed with a HotStart Taq polymerase kit (Qiagen) in a final reaction volume of 25 μL. The reaction mixture consisted of 1× PCR buffer, 0.25 μM of each primer, 0.2 mM dNTPs, 0.0625 U/μL Taq DNA polymerase, and 100 ng DNA template. For the amplification of *tuf*, *als*, *gapC*, *galP*, *gyrB,* and *rpoC* loci, the PCR conditions consisted of an initial denaturation step at 95 °C for 15 min followed by 35 cycles consisting of 94 °C for 45 s, annealing temperatures varying from 56 °C to 58 °C for 45 s, and 72 °C for 1min10s, with a final extension step at 72 °C for 5 min. To amplify the *atpA* locus, the PCR conditions consisted of an initial denaturation step at 95 °C for 15 min, followed by a step of 3 cycles consisting of 95 °C for 1 min, annealing at 56 °C for 2min15s, and 72 °C for 1min15s, another step of 30 cycles consisting of 95 °C for 35 s, annealing at 56 °C for 1min15s, and 72 °C for 1min15s, with a final extension step at 72 °C for 7 min. Primers were purchased from Integrated DNA Technologies (IDT). The cycles were performed in a Veriti 96-Well Thermal Cycler (Thermo Fisher Scientific) and amplicons were separated in QIAxcell Advanced using the QX DNA Screening Kit (Qiagen). The primers used for the MLST study are listed on Supplementary Table S2. Sequencing reactions were performed as described in the previous section.

In order to determine the sequence type (ST) of each isolate, the MLST data generated in this study were compared with the allelic profiles and STs previously published for *L. garvieae* using BLAST from the NCBI website. Arbitrary numbers for distinct allele sequences for each *locus* and new STs were assigned in the order of description following the numerical identification given by Lin et al. (2020) and Thiry et al. (2021). The allele sequences detected at each of the seven loci are listed in Supplementary Table S3. Isolate relationships were analyzed using the eBURST algorithm (Feil et al., 2004). Clonal complexes (CCs) were defined using the SLV bias with default parameters.

### 2.4 Challenge trial and pathogenicity study

To fulfil Koch’s postulates and investigate bacterial virulence, a median lethal dose (LD50) study was performed. After the characterization of the *L. petauri* isolates, one representative isolate (48/20-05) was chosen considering that the ST24 was the most frequently observed among the isolates recovered from all farms. This study was approved by the Animal Experimentation Ethics Committee of Vaxxinova (protocol n. 013/20).

Tilapia fingerlings (average weight 20 g) were obtained from a source with no history of lactococcosis, and a sub-sample (n=10) of the population was confirmed as negative for bacterial infection by complete clinical and bacteriological analysis by culture of the brain and kidney (as previously described) prior to the challenge. Fish were acclimated for seven days in 55 L freshwater tanks at 28 °C and fed commercial tilapia feed three times daily (Nanolis®, ADM). Water quality parameters (temperature, dissolved oxygen, and pH) were monitored and maintained to meet species requirements.

The bacterial isolate was grown in 50 mL BHI broth at 28 °C for 18 h. The initial bacterial suspension was incubated to reach a 0.5 McFarland turbidity standard (∼10^8^ colony-forming units [CFU]/mL), and tenfold serial dilutions (10^6^ to 10^2^ CFU/mL) were prepared in BHI broth. The amount of bacteria on each dilution was calculated by the number of colonies suspended in 0.1 mL BHI broth using the plate count method (Miles et al., 1938). For the challenge, the fish were divided into six experimental groups in duplicate, with 20 fish per group (one tank/treatment). Fish were starved for 24 h prior to the challenge and anesthetized by immersion in clove oil at 50 mg/L before handling. Fish in different treatment groups were challenged with a 0.1 mL intraperitoneal injection of 2.5 × 10^6^, 2.9 × 10^5^, 4.8 × 10^4^, 4.5 × 10^3^, and 4.0 × 10^2^ CFU per fish. Control fish were treated in a similar manner but received sterile BHI broth.

Following the challenge, the fish were monitored for 15 days and mortality was recorded. Fish were collected immediately after death for re-isolation of bacteria. At the end of the trial, all surviving fish were euthanized by immersion in clove oil at 100 mg/L for re-isolation of the bacteria. The re-isolation procedure was performed using the same procedures described previously (section 2.2), using TSA supplemented with 5 % sheep blood. The LD50 was calculated using the method described by Reed and Muench (1938).

### 2.5 Antimicrobial susceptibility analysis

The susceptibility of isolated *L. petauri* to antibiotics was determined using the standard disk diffusion susceptibility test, according to the method described by Bauer et al. (1966), following the Clinical and Laboratory Standard Institute protocols in VET-03 (CLSI, 2020a). For this purpose, antimicrobial disks of amoxicillin (10 μg), erythromycin (15 μg), florfenicol (30 μg), neomycin (10 μg), norfloxacin (10 μg), oxytetracycline (30 μg), and sulfamethoxazole with trimethoprim (25 μg) (Oxoid, Basingstoke, UK) were used. Briefly, *L. petauri* isolates were thawed from freezer stocks and grown on TSA supplemented with 5 % sheep blood at 28 °C for 24 h. Four to six colonies were suspended in sterile saline solution (0.9 %), and absorbance adjusted to match a 0.5 McFarland turbidity standard (optical density of 0.08–0.13 at 625 nm at a concentration of ∼10^8^ CFU/mL). The bacterial suspension was then streaked onto cation-adjusted Mueller Hinton agar (HiMedia) supplemented with 5 % defibrinated sheep blood using a sterile swab. After drying, antimicrobial disks were dispensed onto the surface of the inoculated agar plates and plates were incubated at 28 °C for 24 h, when the diameters of the zones of complete inhibition were measured. *Escherichia coli* ATTC 25922 and *Aeromonas salmonicida* subsp. *salmonicida* ATCC 33658 were used as the quality control isolates.

Since no epidemiological cutoff values (CO_WT_) have been stablished established for *L. petauri* in the literature, provisional CO_WT_ were calculated by the Normalized Resistance Interpretation (NRI) method using automatic and freely available Excel spreadsheet calculators (www.bioscand.se/nri/) (Kronvall, 2003; Kronvall and Smith, 2016). The NRI was used with the permission of the patent holder (Bioscand AB, TABY, Sweden; US Patent No. 7,465,559 and European Patent No. 1383913). The isolates were classified as wild-type (WT) or non-wild-type (NWT) based on the CO_WT_ calculated for each antimicrobial agent.

## 3 Results

### 3.1 Bacterial identification

From the 14 outbreaks examined in this study, a total of 178 isolates were identified as *L. garvieae* by MALDI-TOF mass spectrometry. Subsequently, two isolates were selected from each outbreak, and the expected amplicons for the 16S rRNA (∼1500 bp) were obtained from these isolates and compared to the NCBI database. However, it was not possible to determine the species of the isolates from this analysis because the 16S rRNA gene similarities were higher than 99 % for *L. petauri* and *L. garvieae* (data not shown).

Consequently, a phylogenetic tree based on the *gyrB* gene sequence (1,340 bp) was constructed (Figure 3). Most isolates were closely related to *L. petauri* strains B1726, PAQ102015-99, CF11, and 159469. Therefore, these isolates were classified as *L. petauri*. Two isolates (30/19-01 and 05/21-28) were closely related to *L. garvieae* strains ATCC49156, Lg2, JJJN1. Accordingly, these two isolates were classified as *L. garvieae*.

**Figure 3.**
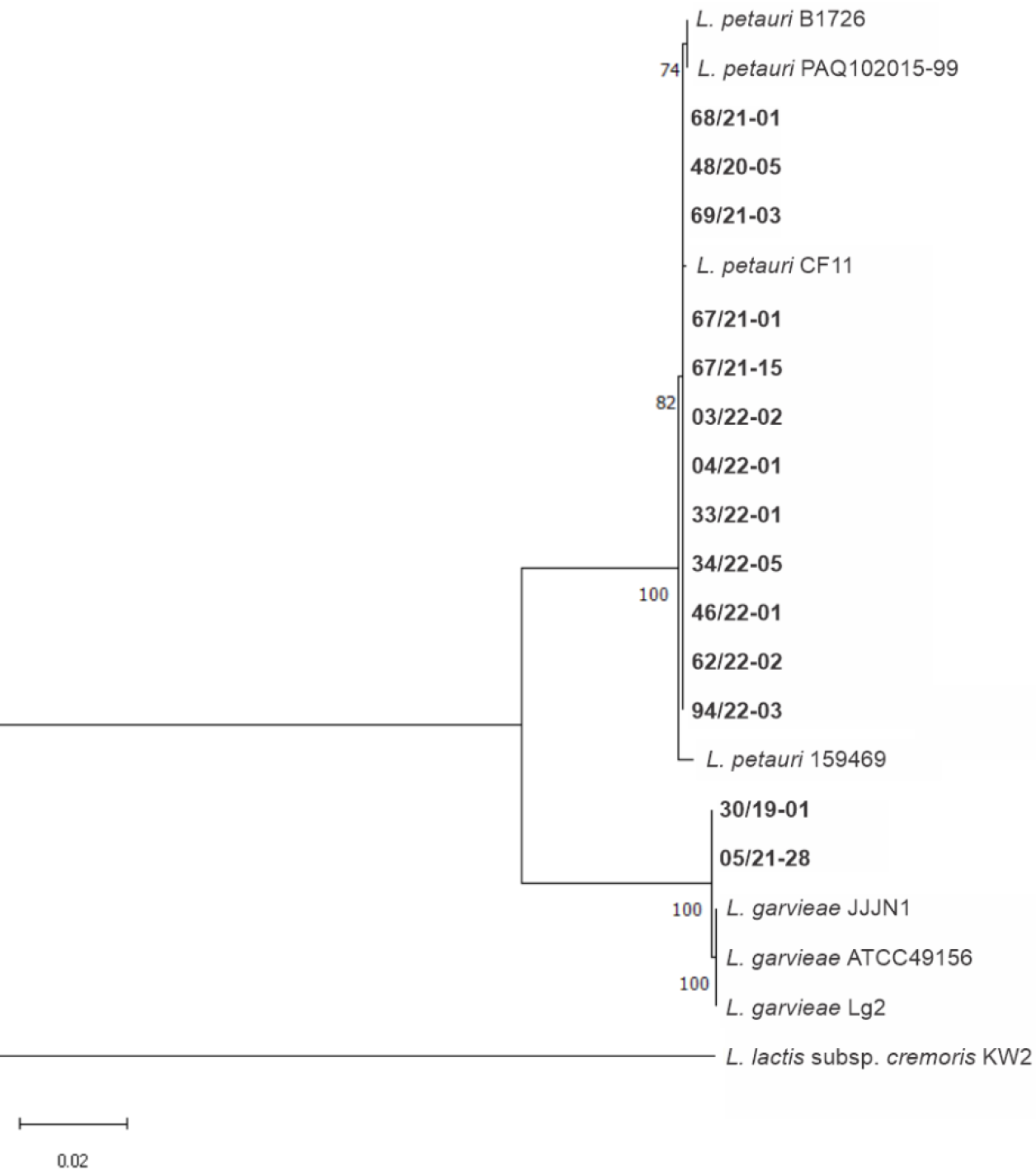
Neighbor-joining phylogenetic tree based on the *gyrB* gene sequences (1,340 bp) of the isolates from this study (in bold) relative to other representative strains within the species *L. petauri* and *L. garvieae* (bootstrap test, 1,000 replicates). *Lactococcus lactis* subsp. *cremoris* KW2 is presented as the outgroup.

Furthermore, the *gyrB* gene sequences from *L. petauri* isolates obtained in this study presented similarities higher than 99.63 % with *L. petauri* reference strains (B1726, PAQ102015-99, CF11, and 159469). Conversely, these isolates presented 93.66 % similarity when compared to the *gyrB* gene sequences from *L. garvieae* reference strains (ATCC49156, Lg2, JJJN1) and 93.73 % similarity with *L. garvieae* isolates from this study. Likewise, the *gyrB* gene sequences from *L. garvieae* isolates obtained in this study presented 99.93 % similarity with *L. garvieae* reference strains (ATCC49156, Lg2, JJJN1) and 93.66 % similarity with *L. petauri* reference strains (B1726, PAQ102015-99, CF11, and 159469).

During the monitoring of outbreaks, coinfection with other bacteria (*Aeromonas* spp., *Edwardsiella* spp., *S. agalactiae*, and *S. iniae*) was identified by means of MALDI TOF mass spectrometry in some fish from four farms (Table 1). Infections by *Aeromonas* spp., *Edwardsiella* spp., *Francisella orientalis*, *S. agalactiae*, *S. dysgalactiae, S. iniae*, and *Vibrio vulnificus* were also diagnosed in sampled fish, but not as coinfection.

### 3.2 MLST and eBURST profiles

The 32 isolates analyzed were grouped in three different STs, one previously reported for *L. garvieae* isolates (ST24) and two new STs, ST46 and ST47 (Table 2). The ST46 presented a new allele for *gyrB*, which was characterized as allele 27 for this gene. The new allele *gyrB* 27 presented a single nucleotide polymorphism (SNP) at position 677 in relation to the allele *gyrB* 10, in which a cytosine (C) was replaced by a thymine (T). The ST47 presented a new allele for *als*, which was characterized as allele 29 for this gene, and a new allele for *atpA*, which was characterized as allele 21 for this gene. The new allele *als* 29 presented 11 SNPs in relation to the allele *als* 10: at position 7, a guanine (G) was replaced by an adenine (A); at 226, a T by a C; at 229, a T by a G; at 250, a C by a T; at 283, a T by an A; at 298, an A by a G; at 355, a T by a C; at 409, an A by a G; at 436, a C by a T; at 535, a T by a C; and at 622, an A by a G. The new allele *atpA* 21 presented one SNP in relation to the allele *aptA* 6, in which a C was replaced by a T at position 342.

**Table 2.** Alelic profiles, sequence types (ST), and clonal complexes (CCs) of *L. Petauri* and *L. garvieae* isolates

| Isolates | Species | Alelic profile |  |  |  |  |  |  | ST | CC |
| --- | --- | --- | --- | --- | --- | --- | --- | --- | --- | --- |
|  |  | *als* | *galP* | *gapC* | *gyrB* | *atpA* | *rpoC* | *tuf* |  |  |
| 48/20-02 | *L. petauri* | 17 | 16 | 2 | 7 | 12 | 9 | 3 | 24 | Singleton |
| 48/20-05 | *L. petauri* | 17 | 16 | 2 | 7 | 12 | 9 | 3 | 24 | Singleton |
| 48/20-08 | *L. petauri* | 17 | 16 | 2 | 7 | 12 | 9 | 3 | 24 | Singleton |
| 48/20-09 | *L. petauri* | 17 | 16 | 2 | 7 | 12 | 9 | 3 | 24 | Singleton |
| 48/20-10 | *L. petauri* | 17 | 16 | 2 | 7 | 12 | 9 | 3 | 24 | Singleton |
| 48/20-11 | *L. petauri* | 17 | 16 | 2 | 7 | 12 | 9 | 3 | 24 | Singleton |
| 48/20-12 | *L. petauri* | 17 | 16 | 2 | 7 | 12 | 9 | 3 | 24 | Singleton |
| 48/20-14 | *L. petauri* | 17 | 16 | 2 | 7 | 12 | 9 | 3 | 24 | Singleton |
| 48/20-16 | *L. petauri* | 17 | 16 | 2 | 7 | 12 | 9 | 3 | 24 | Singleton |
| 30/19-01 | *L. garvieae* | 12 | 12 | 7 | 27 | 8 | 13 | 6 | 46 | 17 |
| 05/21-28 | *L. garvieae* | 12 | 12 | 7 | 27 | 8 | 13 | 6 | 46 | 17 |
| 67/21-01 | *L. petauri* | 17 | 16 | 2 | 7 | 12 | 9 | 3 | 24 | Singleton |
| 67/21-05 | *L. petauri* | 17 | 16 | 2 | 7 | 12 | 9 | 3 | 24 | Singleton |
| 68/21-01 | *L. petauri* | 17 | 16 | 2 | 7 | 12 | 9 | 3 | 24 | Singleton |
| 68/21-05 | *L. petauri* | 17 | 16 | 2 | 7 | 12 | 9 | 3 | 24 | Singleton |
| 69/21-03 | *L. petauri* | 17 | 16 | 2 | 7 | 12 | 9 | 3 | 24 | Singleton |
| 69/21-07 | *L. petauri* | 17 | 16 | 2 | 7 | 12 | 9 | 3 | 24 | Singleton |
| 67/21-15 | *L. petauri* | 17 | 16 | 2 | 7 | 12 | 9 | 3 | 24 | Singleton |
| 67/21-18 | *L. petauri* | 17 | 16 | 2 | 7 | 12 | 9 | 3 | 24 | Singleton |
| 03/22-02 | *L. petauri* | 17 | 16 | 2 | 7 | 12 | 9 | 3 | 24 | Singleton |
| 03/22-09 | *L. petauri* | 17 | 16 | 2 | 7 | 12 | 9 | 3 | 24 | Singleton |
| 04/22-01 | *L. petauri* | 17 | 16 | 2 | 7 | 12 | 9 | 3 | 24 | Singleton |
| 04/22-07 | *L. petauri* | 17 | 16 | 2 | 7 | 12 | 9 | 3 | 24 | Singleton |
| 33/22-01 | *L. petauri* | 17 | 16 | 2 | 7 | 12 | 9 | 3 | 24 | Singleton |
| 33/22-03 | *L. petauri* | 17 | 16 | 2 | 7 | 12 | 9 | 3 | 24 | Singleton |
| 34/22-05 | *L. petauri* | 17 | 16 | 2 | 7 | 12 | 9 | 3 | 24 | Singleton |
| 34/22-06 | *L. petauri* | 17 | 16 | 2 | 7 | 12 | 9 | 3 | 24 | Singleton |
| 46/22-01 | *L. petauri* | 17 | 16 | 2 | 7 | 12 | 9 | 3 | 24 | Singleton |
| 46/22-05 | *L. petauri* | 17 | 16 | 2 | 7 | 12 | 9 | 3 | 24 | Singleton |
| 62/22-02 | *L. petauri* | 17 | 16 | 2 | 7 | 12 | 9 | 3 | 24 | Singleton |
| 62/22-04 | *L. petauri* | 17 | 16 | 2 | 7 | 12 | 9 | 3 | 24 | Singleton |
| 94/22-03 | *L. petauri* | 29 | 6 | 2 | 7 | 21 | 11 | 7 | 47 | Singleton |

Most *L. petauri* isolates were grouped in ST24, whereas one isolate belonged to ST47. Regarding the *L. garvieae* isolates identified, both were grouped in ST46.

The eBURST analysis from MLST was performed to evaluate the evolutionary relationships between *L. petauri* and *L. garvieae* STs (Figure 4). The ST46 was grouped into CC17 with ST16 and ST17, with ST17 as the founder. Whereas the ST24 and ST47 were characterized as singletons.

**Figure 4.**
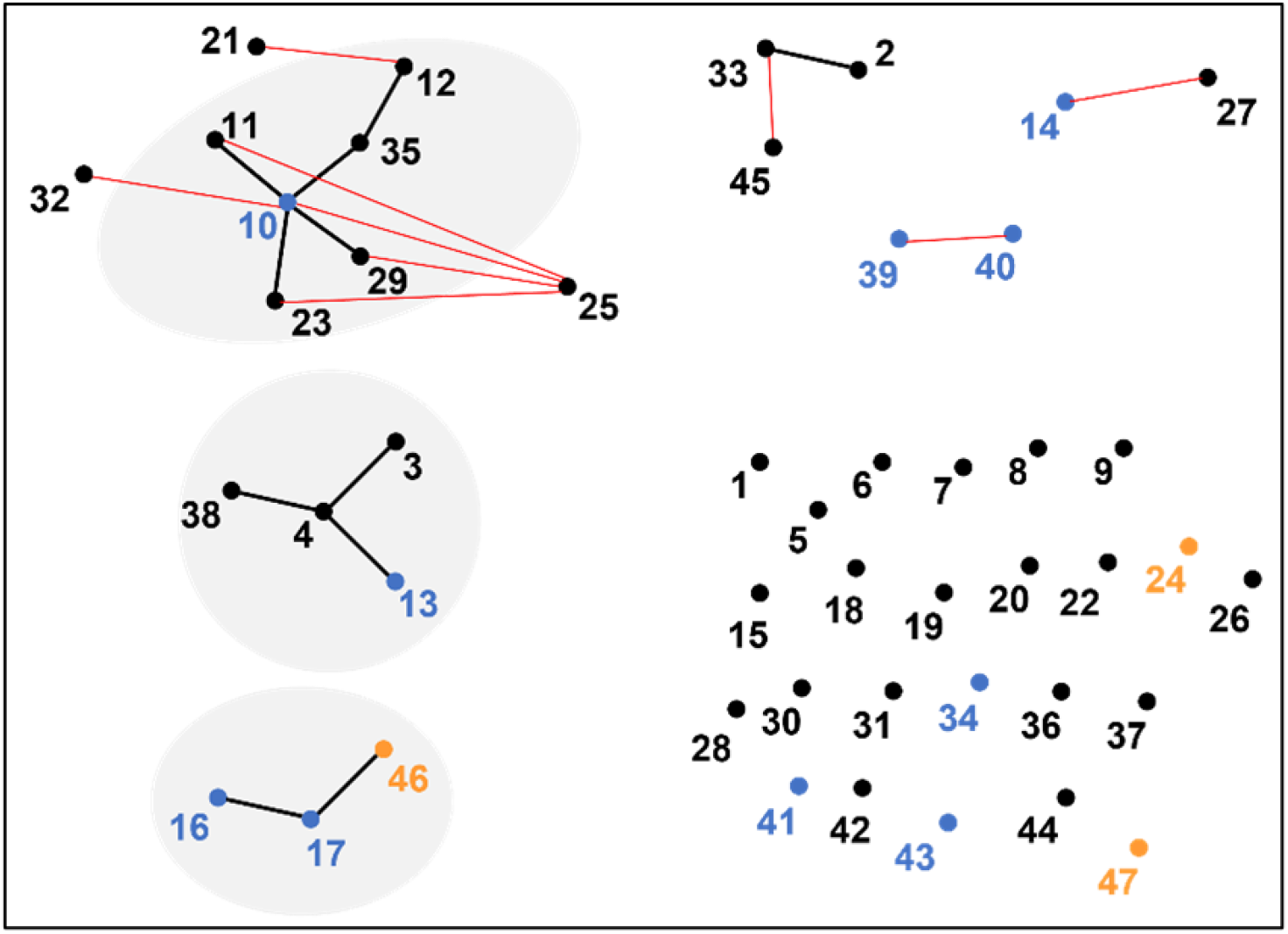
Evolutionary relationships between *L. petauri* and *L. garvieae* through eBURST analysis from *L. garvieae* MLST scheme. Points represent sequence types (STs), blue points represent STs isolated from fish, orange points represent STs observed in this study, gray circles represent clonal complexes (CCs), black lines represent single-locus variants, red lines represent double-loci variants, and isolated points represent singletons. eBurst group CC17 is formed by strains isolated from fish (Nile tilapia and yellowtail) and human feces.

### 3.3 Challenge trial and pathogenicity study

Nile tilapia challenged with *L. petauri* developed clinical signs within 48 h of challenge. Experimentally infected fish exhibited hyporexia, which was first observed in fish from groups challenged with 2.9 × 10^5^ and 2.5 × 10^6^ CFU/fish. On the following days, fish from the other challenged groups also exhibited hyporexia. The first mortalities were recorded within 48 h in the group of fish challenged with 2.9 × 10^5^ CFU/fish. All groups of experimentally infected fish presented mortalities by 4 days post-challenge (dpc). Mortalities of challenged fish occurred every day until 9 dpc. On the same day, fish from all challenged groups also recovered from hyporexia and exhibited normal feeding behavior until the end of the experimental trial. New mortalities occurred at 13 and 14 dpc in groups challenged with 2.9 × 10^5^ and 4.8 × 10^4^ CFU/fish, respectively (Figure 5). The estimated LD50 for the *L. petauri* isolate used in the challenge was 5.74 × 10^3^ CFU 15 dpc. *L. petauri* was reisolated from the brain and posterior kidney of all dead fish, but it was not reisolated from the surviving fish. No mortalities were observed in the control group.

**Figure 5.**
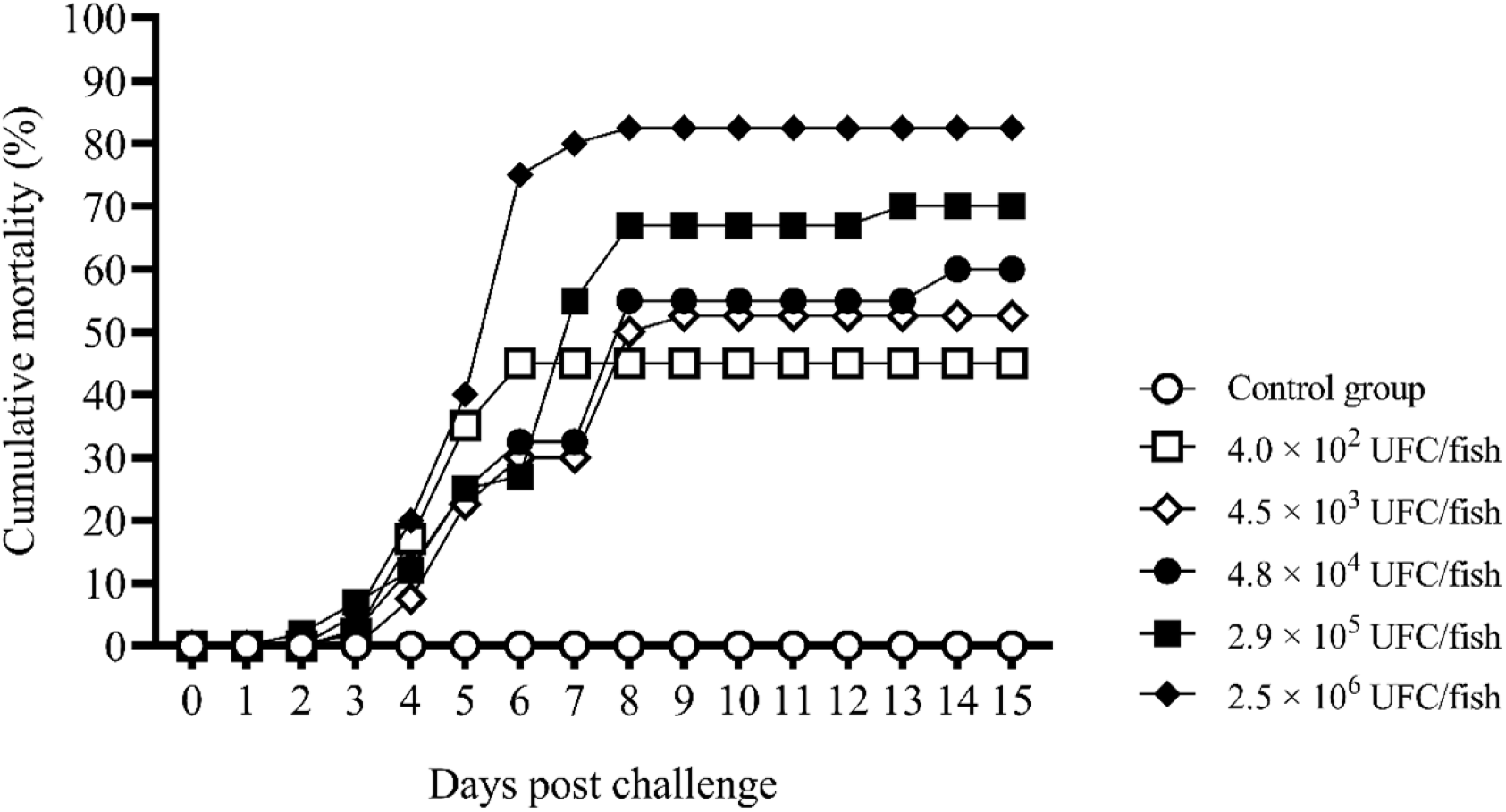
Cumulative percentage mortalities of Nile tilapia challenged with different doses of *L. petauri* by intraperitoneal injection.

### 3.5 Antimicrobial susceptibility analysis

The reference strains *E. coli* ATTC 25922 and *A. salmonicida* subsp. *salmonicida* ATCC 33658 profiles were within ranges established by CLSI (CLSI, 2020b). The calculated provisional CO_WT_ for the antimicrobials and the diameters of the zones of complete inhibition of the antimicrobials against *L. petauri* isolates are presented in Table 3. Except for one isolate (48/20-14), all isolates presented a zone of complete inhibition of 6 mm for sulfamethoxazole-trimethoprim, which prevented the establishment of the CO_WT_ for this antimicrobial. All the isolates were characterized as WT for neomycin and oxytetracycline. In addition, 96.67 % of the isolates were characterized as WT for amoxicillin, erythromycin, and florfenicol, whereas 83.33 % were WT for norfloxacin.

**Table 3.** Provisional epidemiological cutoff values (CO_WT_) and diameters (mm) of the zones of complete inhibition of antimicrobial agents against *L. petauri* isolates determined using standard disk diffusion susceptibility test. Isolates were classified as wild-type (WT) or non-wild-type (NWT) following the provisional CO_WT_ established by the Normalized Resistance Interpretation (NRI) method. Reference strains *E. coli* ATTC 25922 and *A. salmonicida* subsp. *salmonicida* ATCC 33658 within ranges (CLSI, 2020b).

| Isolates | Antimicrobial agents (CO<sub>WT</sub>) |  |  |  |  |  | SXT |
| --- | --- | --- | --- | --- | --- | --- | --- |
|  | AM<br>(16 mm) | ERY<br>(23 mm) | FFC<br>(22 mm) | NEO<br>(9 mm) | NOR<br>(13 mm) | OXT<br>(21 mm) |  |
| 48/20-02 | 20 (WT) | 25 (WT) | 26 (WT) | 11 (WT) | 16 (WT) | 23 (WT) | 6 |
| 48/20-05 | 21 (WT) | 21 (NWT) | 21 (NWT) | 13 (WT) | 13 (WT) | 27 (WT) | 6 |
| 48/20-08 | 34 (WT) | 27 (WT) | 24 (WT) | 12 (WT) | 20 (WT) | 27 (WT) | 6 |
| 48/20-09 | 25 (WT) | 25 (WT) | 25 (WT) | 12 (WT) | 17 (WT) | 25 (WT) | 6 |
| 48/20-10 | 24 (WT) | 25 (WT) | 25 (WT) | 12 (WT) | 17 (WT) | 27 (WT) | 6 |
| 48/20-11 | 24 (WT) | 26 (WT) | 25 (WT) | 11 (WT) | 16 (WT) | 27 (WT) | 6 |
| 48/20-12 | 22 (WT) | 25 (WT) | 25 (WT) | 13 (WT) | 18 (WT) | 26 (WT) | 6 |
| 48/20-14 | 37 (WT) | 34 (WT) | 33 (WT) | 15 (WT) | 21 (WT) | 37 (WT) | 14 |
| 48/20-16 | 23 (WT) | 25 (WT) | 25 (WT) | 11 (WT) | 16 (WT) | 27 (WT) | 6 |
| 67/21-01 | 25 (WT) | 27 (WT) | 27 (WT) | 13 (WT) | 18 (WT) | 27 (WT) | 6 |
| 67/21-05 | 22 (WT) | 25 (WT) | 26 (WT) | 12 (WT) | 13 (WT) | 26 (WT) | 6 |
| 68/21-01 | 24 (WT) | 27 (WT) | 28 (WT) | 13 (WT) | 18 (WT) | 27 (WT) | 6 |
| 68/21-05 | 24 (WT) | 25 (WT) | 25 (WT) | 13 (WT) | 16 (WT) | 27 (WT) | 6 |
| 69/21-03 | 24 (WT) | 26 (WT) | 25 (WT) | 12 (WT) | 17 (WT) | 26 (WT) | 6 |
| 69/21-07 | 25 (WT) | 26 (WT) | 26 (WT) | 13 (WT) | 18 (WT) | 28 (WT) | 6 |
| 67/21-15 | 24 (WT) | 27 (WT) | 26 (WT) | 13 (WT) | 18 (WT) | 29 (WT) | 6 |
| 67/21-18 | 24 (WT) | 28 (WT) | 27 (WT) | 12 (WT) | 19 (WT) | 29 (WT) | 6 |
| 03/22-02 | 28 (WT) | 26 (WT) | 24 (WT) | 12 (WT) | 16 (WT) | 25 (WT) | 6 |
| 03/22-09 | 20 (WT) | 26 (WT) | 25 (WT) | 13 (WT) | 17 (WT) | 27 (WT) | 6 |
| 04/22-01 | 12 (NWT) | 27 (WT) | 26 (WT) | 13 (WT) | 12 (NWT) | 25 (WT) | 6 |
| 04/22-07 | 24 (WT) | 26 (WT) | 26 (WT) | 12 (WT) | 16 (WT) | 26 (WT) | 6 |
| 33/22-01 | 27 (WT) | 25 (WT) | 23 (WT) | 13 (WT) | 16 (WT) | 26 (WT) | 6 |
| 33/22-03 | 23 (WT) | 25 (WT) | 25 (WT) | 13 (WT) | 11 (NWT) | 26 (WT) | 6 |
| 34/22-05 | 24 (WT) | 25 (WT) | 25 (WT) | 14 (WT) | 18 (WT) | 26 (WT) | 6 |
| 34/22-06 | 24 (WT) | 25 (WT) | 25 (WT) | 13 (WT) | 17 (WT) | 26 (WT) | 6 |
| 46/22-01 | 25 (WT) | 26 (WT) | 26 (WT) | 13 (WT) | 11 (NWT) | 28 (WT) | 6 |
| 46/22-05 | 26 (WT) | 27 (WT) | 27 (WT) | 13 (WT) | 12 (NWT) | 26 (WT) | 6 |
| 62/22-02 | 23 (WT) | 26 (WT) | 27 (WT) | 13 (WT) | 16 (WT) | 27 (WT) | 6 |
| 62/22-04 | 22 (WT) | 25 (WT) | 26 (WT) | 12 (WT) | 13 (WT) | 26 (WT) | 6 |
| 94/22-03 | 25 (WT) | 27 (WT) | 26 (WT) | 15 (WT) | 11 (NWT) | 30 (WT) | 6 |

Abbreviations: AM: amoxicillin, ERY: erythromycin, FFC: florfenicol, NEO:
neomycin, NOR: norfloxacin, OXT: oxytetracycline, SXT: sulfamethoxazole with
trimethoprim.
Grey colored cells indicate isolates classified as NWT for that antimicrobial agent.

## 4 Discussion

Considering that *L. petauri* is a recently described species of the genus *Lactococcus*, phylogenetic analyses were performed in this study to correctly determine the taxonomic classification of the isolates obtained from diseased Nile tilapia in Brazil. Phylogenetic analysis based on the *gyrB* gene sequence revealed that 30 out of 32 isolates belonged to the species *L. petauri*, whereas the other two isolates belonged to *L. garvieae*. The *gyrB* gene sequence has been used combined with multilocus sequence analysis (MLSA) to differentiate between *L. petauri* and *L. garvieae* (Shahin et al., 2022). In the present study, the *gyrB* gene sequence also allowed the differentiation between the two species, since isolates from these species were grouped in distinct branches of the phylogenetic tree.

On the other hand, 16S rRNA gene analysis did not provide information that allowed to determine the species to which the isolates from this study belonged. This corroborates the literature, since it has been previously reported that the analysis of the 16s rRNA gene is not sufficient to discriminate between *L. petauri* and *L. garvieae* (Ou et al., 2020). The initial identification of the bacterial species using MALDI-TOF mass spectrometry also did not differentiate between the two species, and all isolates were initially classified as *L. garvieae*. This was expected, as *L. petauri* is not currently recognized by the microbial peptide mass spectrum database.

Different approaches have been used to assess *L. garvieae* genetic diversity, including enterobacterial repetitive intergenic consensus-PCR (ERIC-PCR), repetitive element palindromic-PCR (REP-PCR), and MLST (Feito et al., 2022; Ortega et al., 2020; Reguera-Brito et al., 2016; Torres-Corral and Santos, 2021). Although the literature reveals that *L. garvieae* is a bacterial species with high genetic heterogeneity, fish isolates recovered from the same outbreak generally present a clonal structure. Considering that MLST is a well-established methodology for the study of *L. garvieae* genetic diversity (Ferrario et al., 2013; Lin et al., 2020; Reguera-Brito et al., 2016) and that *L. petauri* and *L. garvieae* share high genetic similarity (Goodman et al., 2017; Kotzamanidis et al., 2020; Martinovic et al., 2021; Ou et al., 2020), the MLST scheme for *L. garvieae* was used in this study to analyze *L. petauri* isolates. In addition, the MLST analysis allows the understanding of evolutionary stories between bacterial lineages and epidemiological tracking of bacteria (Maiden et al., 2013; Pérez-Losada et al., 2013). The MLST scheme for *L. garvieae* has proved to be a useful tool for understanding the molecular epidemiology of *L. petauri*. Therefore, the establishment of a PubMLST for *L. petauri* and *L. garvieae* should be encouraged to unify and consolidate these valuable data for the two species.

The studied isolates could be classified into three STs (lineages), including two newly described STs. Isolates of *L. petauri* presented two STs, ST24 and ST47, which were characterized as singletons. Previously, ST24 has been described in samples from human feces and human gall bladder (Lin et al., 2020), whereas ST47 is a newly described ST. In this study, *L. petauri* ST24 was the most common lineage observed. This might indicate that this lineage is well adapted to infect Nile tilapia. Moreover, ST24 did not present relationships with other STs, which further indicated this adaptation. The *L. garvieae* ST46 is also a newly described ST, and it forms a CC with ST16, which was previously identified in yellowtail, and ST17, which was identified in yellowtail and human feces (Lin et al., 2020).

The three lineages presented a geographical distribution pattern, with ST24 occurring mainly in the northeast region of Brazil and the ST46 and ST47 occurring exclusively in the Southeast. The spatial distribution of *L. petauri* and *L. garvieae* demonstrated that these pathogens have spread throughout Brazil. This raises the hypothesis of pathogen transmission by animal movement, probably through the fingerling trade. Furthermore, *L. petauri* ST24 was observed in different regions of the country, which illustrates a rapid expansion of this *L. petauri* lineage.

To the best of our knowledge, *L. petauri* has not been described infecting Nile tilapia previously. The most important pathogens that have been identified to cause disease in Nile tilapia are bacterial agents, such as *Streptococcus* spp., *F. orientalis*, *Flavobacterium columnare*, *Edwardsiella* spp. (Armwood et al., 2019; Barony et al., 2017, 2015; Delphino et al., 2019; Evans et al., 2015; Figueiredo et al., 2012; Leal et al., 2014; Reichley et al., 2017), as well as viral agents, such as tilapia lake virus (TILV) and infectious spleen and kidney necrosis virus (ISKNV) (Eyngor et al., 2014; Subramaniam et al., 2016).

Infections of *L. garvieae* in Nile tilapia have been more frequently reported in the last years (Anshary et al., 2014; Osman et al., 2017; Tsai et al., 2012). Although *L. garvieae* has been reported in Nile tilapia in Brazil (Evans et al., 2009; Sebastião et al., 2015), the identification of this this pathogen was not frequent in routine bacteriological diagnosis in fish until 2019. Although this bacterium has been associated with clinical signs in infected Nile tilapia, it has not been frequently linked to high mortality rates in this fish. Nevertheless, *L. garvieae* outbreaks in Nile tilapia in Zambia in 2015/16 (Bwalya et al., 2020b) raised an alarm regarding the potential impact of this pathogen on tilapia aquaculture. Therefore, during the first outbreak identified in 2020, when 26 out of 66 sampled fish were infected with *L. petauri*, it was clear that greater attention should be given to this pathogen. Then, other outbreaks followed, which reinforced the importance of *L. petauri* in Nile tilapia.

In some of the investigated outbreaks, more than half of the sampled fish were infected by *L. petauri*, whereas in other outbreaks the infection was detected in only one or three of the sampled fish. This was probably due to the different stages of the disease in these systems. Therefore, while some outbreaks were detected when the disease had already spread on the farm, in others the disease was either starting to spread or it had already been overcome by the fish. Additionally, this pathogen was repeatedly isolated in farm 4, what indicates the maintenance of *L. petauri* for long periods in the farm production system.

Interestingly, coinfections occurred in five of the 14 outbreaks studied, whereas concurrent infections in the same farm were identified in all outbreaks. This indicates that bacterial coinfections probably do not play a significant role in the dynamics of *L. petauri* infection in Nile tilapia. On the other hand, the temperature seemes to be critical for the disease. Most of the studied outbreaks occurred during the warmer months (November to March) at temperatures above 26 °C. One exception occurred on Farm 12, where the *L. petauri* ST47 was identified in one out of 22 sampled fish in June, when the temperature was 22 °C. Not surprisingly, in the outbreaks that *L. garvieae* was identified, the temperatures (above 25.6 °C) were similar to those registered during *L. petauri* outbreaks. This agrees with what has been reported for *L. petauri* outbreaks in rainbow trout (Kotzamanidis et al., 2020) and *L. garvieae* outbreaks in Nile tilapia, rainbow trout, and cobia (Bwalya et al., 2020b; Ortega et al., 2020; Rao et al., 2021), which occurred at higher temperatures.

In the challenge trial, Nile tilapia infected with *L. petauri* developed hyporexia, whereas none of the signs reported by the producers during the outbreaks were observed in the experimental trial. Other authors have described that Nile tilapia experimentally infected with *L. garvieae* exhibited darkening of the skin, lethargy, exophthalmia and corneal opacity (Bwalya et al., 2020a, 2020b). The deaths peak occurred between 3–8 dpc, which demonstrated that *L. petauri* infection in Nile tilapia progressed very rapidly. The estimated LD50 of 5.74 × 10^3^ CFU 15 dpc was lower than previously estimated values of LD50 for *L. garvieae* in Nile tilapia of 9.6 × 10^5^ CFU and 1.41 × 10^5^ CFU using the same infection route (Bwalya et al., 2021; Evans et al., 2009). This might indicate that *L. petauri* is more virulent than *L. garvieae* to Nile tilapia. However, this difference may also be related to fish susceptibility which is influenced by age, size, and bacterial strain, among others.

Another intriguing finding from the challenge trial was that *L. petauri* was not reisolated from the surviving fish at 15 dpc. Similar findings have been reported for *L. garvieae* infection in Nile tilapia by Bwalya et al. (2020a). Therefore, it can be hypothesized that Nile tilapia that do not succumb to lactococcosis within this time may recover from the infection and eliminate the pathogen from their organism. The pathogenicity of *L. garvieae* isolates obtained in this study was not assessed since this bacterium has already been proven to be pathogenic to Nile tilapia (Anshary et al., 2014; Baba et al., 2017; Bwalya et al., 2020b; Evans et al., 2009).

Regarding antimicrobial resistance, most of the *L. petauri* isolates were classified as WT to the antimicrobials analyzed. Another study on *L. petauri* isolated from rainbow trout reported that the isolates were sensitive to erythromycin and florfenicol, but resistant to tetracycline (Shahin et al., 2021). However, different methods have been used to determine antimicrobial, making it difficult to compare the data. The literature concerning *L. garvieae* antimicrobial resistance profiles shows great heterogeneity, probably as a result of the misuse or overuse of antimicrobial agents at the farm level. In addition, the lack of established specific susceptibility breakpoints for *L. garvieae* contributes to these variations, as many studies have used the breakpoints recommended for streptococci (Gibello et al., 2016). In this study, provisional CO_WT_ were determined for *L. petauri* recovered from fish for seven antimicrobial agents from different drug classes. The establishment of susceptibility breakpoints for *L. petauri* may improve the routine use of antimicrobial susceptibility testing for this pathogen, thus allowing appropriate drug selection in fish farming and monitoring of antimicrobial resistance among isolates and outbreaks. Nevertheless, the CO_WT_ calculated in this study are provisional and may vary according to the geographic regions of the bacterial isolates. Therefore, antimicrobial resistance analysis of a larger number of *L. petauri* isolates from different locations can contribute to the establishment of a CO_WT_.

Some isolates recovered from the same farm exhibited different antimicrobial resistance phenotypes. Although the isolates were genetically similar according to MLST (same ST lineage), some genetic differences among isolates may contribute to this phenotypic variation. The presence of antimicrobial resistance genes in the genome of *L. Petauri* has been previously described (Kotzamanidis et al., 2020; Ou et al., 2020) and will be further assessed in the genome of some isolates from this study for comparison with the phenotypic profile of the isolates.

## 5 Conclusions

This study described *L. petauri* infection for the first time in Nile tilapia. A total of 14 lactococcosis outbreaks were analyzed from 2019 to 2022, of which 12 had been caused by *L. petauri* and two by *L. garvieae*. Furthermore, *L. petauri* pathogenicity to Nile tilapia was confirmed trough Koch’s postulates. Analysis of *gyrB* gene sequence can be used to differentiate *L. petauri* from *L. garvieae*, thus allowing the correct identification of these emerging fish pathogens. Two MLST lineages of *L. petauri* were observed: ST24, previously described as *L. garvieae*, and the new ST47. In the present study, *L. petauri* was observed in different Brazilian regions, illustrating a rapid expansion of this pathogen. Also, most *L. petauri* isolates exhibited susceptible profiles to the analyzed antimicrobials.

## Funding

This work was supported by the Brazilian fostering agencies Coordenação de Aperfeiçoamento de Pessoal de Nível Superior (CAPES); Fundação de Amparo à Pesquisa do Estado de Minas Gerais (FAPEMIG); and Conselho Nacional de Desenvolvimento Científico e Tecnológico (CNPq) [Grant number 315995/2021-1].

## Notes

### Competing Interest Statement

The authors have declared no competing interest.

## References

Anshary, H., Kurniawan, R.A., Sriwulan, S., Ramli, R., Baxa, D. v., 2014. Isolation and molecular identification of the etiological agents of streptococcosis in Nile tilapia (*Oreochromis niloticus*) cultured in net cages in Lake Sentani, Papua, Indonesia. Springerplus 3. https://doi.org/10.1186/2193-1801-3-627

Armwood, A.R., Camus, A.C., López-Porras, A., Ware, C., Griffin, M.J., Soto, E., 2019. Pathologic changes in cultured Nile tilapia (*Oreochromis niloticus*) associated with an outbreak of *Edwardsiella anguillarum*. Journal of Fish Diseases 42, 1463–1469. https://doi.org/10.1111/jfd.13058

Assis, G.B.N., Pereira, F.L., Zegarra, A.U., Tavares, G.C., Leal, C.A., Figueiredo, H.C.P., 2017. Use of MALDI-TOF mass spectrometry for the fast identification of gram-positive fish pathogens. Frontiers in Microbiology 8. https://doi.org/10.3389/fmicb.2017.01492

Baba, E., Acar, Ü., Yılmaz, S., Öntaş, C., Kesbiç, O.S., 2017. Pre-challenge and post-challenge haemato-immunological changes in *Oreochromis niloticus* (Linnaeus, 1758) fed argan oil against *Lactococcus garvieae*. Aquaculture Research 48, 4563–4572. https://doi.org/10.1111/are.13282

Barony, G.M., Tavares, G.C., Assis, G.B.N., Luz, R.K., Figueiredo, H.C.P., Leal, C.A.G., 2015. New hosts and genetic diversity of *Flavobacterium columnare* isolated from Brazilian native species and Nile tilapia. Diseases of Aquatic Organisms 117, 1–11. https://doi.org/10.3354/dao02931

Barony, G.M., Tavares, G.C., Pereira, F.L., Carvalho, A.F., Dorella, F.A., Leal, C.A.G., Figueiredo, H.C.P., 2017. Large-scale genomic analyses reveal the population structure and evolutionary trends of *Streptococcus agalactiae* strains in Brazilian fish farms. Scientific Reports 7, 1–10. https://doi.org/10.1038/s41598-017-13228-z

Bauer, A.W., Kirby, W.M., Sherris, J.C., Turck, M., 1966. Antibiotic susceptibility testing by a standardized single disk method. Am J Clin Pathol 45, 493–6.

Bwalya, P., Hang’ombe, B.M., Evensen, Ø., Mutoloki, S., 2021. *Lactococcus garvieae* isolated from Lake Kariba (Zambia) has low invasive potential in Nile tilapia (*Oreochromis niloticus*). Journal of Fish Diseases 44, 721–727. https://doi.org/10.1111/jfd.13339

Bwalya, P., Hang’Ombe, B.M., Gamil, A.A., Munang’Andu, H.M., Evensen, Ø., Mutoloki, S., 2020a. A whole-cell *Lactococcus garvieae* autovaccine protects Nile tilapia against infection. PLoS ONE 15. https://doi.org/10.1371/journal.pone.0230739

Bwalya, P., Simukoko, C., Hang’ombe, B.M., Støre, S.C., Støre, P., Gamil, A.A.A., Evensen, Ø., Mutoloki, S., 2020b. Characterization of streptococcus-like bacteria from diseased Oreochromis niloticus farmed on Lake Kariba in Zambia. Aquaculture 523. https://doi.org/10.1016/j.aquaculture.2020.735185

Chan, J.F.W., Woo, P.C.Y., Teng, J.L.L., Lau, S.K.P., Leung, S.S.M., Tam, F.C.C., Yuen, K.Y., 2011. Primary infective spondylodiscitis caused by *Lactococcus garvieae* and a review of human *L. garvieae* infections. Infection. https://doi.org/10.1007/s15010-011-0094-8

CLSI, 2020a. Methods for Antimicrobial Broth Dilution and Disk Diffusion Susceptibility Testing of Bacteria Isolated From Aquatic Animals.CLSI guideline VET03, 2nd ed. ed. Clinical and Laboratory Standards Institute, Wayne, PA.

CLSI, 2020b. Performance Standards for Antimicrobial Susceptibility Testing of Bacteria Isolated From Aquatic Animals. CLSI supplement VET04, 3rd ed. ed. Clinical and Laboratory Standards Institute, Wayne, PA.

Delphino, M.K.V.C., Leal, C.A.G., Gardner, I.A., Assis, G.B.N., Roriz, G.D., Ferreira, F., Figueiredo, H.C.P., Gonçalves, V.S.P., 2019. Seasonal dynamics of bacterial pathogens of Nile tilapia farmed in a Brazilian reservoir. Aquaculture 498, 100–108. https://doi.org/10.1016/j.aquaculture.2018.08.023

Evans, J.J., Klesius, P.H., Shoemaker, C.A., 2009. First isolation and characterization of *Lactococcus garvieae* from Brazilian Nile tilapia, *Oreochromis niloticus* (L.), and pintado, *Pseudoplathystoma corruscans* (Spix & Agassiz). Journal of Fish Diseases 32, 943–951. https://doi.org/10.1111/j.1365-2761.2009.01075.x

Evans, J.J., Pasnik, D.J., Klesius, P.H., 2015. Differential pathogenicity of five *Streptococcus agalactiae* isolates of diverse geographic origin in Nile tilapia (*Oreochromis niloticus* L.). Aquaculture Research 46, 2374–2381. https://doi.org/10.1111/are.12393

Evans, J.J., Pasnik, D.J., Klesius, P.H., Al-Ablani, S., 2006. First report of *Streptococcus agalactiae* and *Lactococcus garvieae* from a wild bottlenose dolphin (*Tursiops truncatus*). Journal of Wildlife Diseases 42, 561–569.

Eyngor, M., Zamostiano, R., Tsofack, J.E.K., Berkowitz, A., Bercovier, H., Tinman, S., Lev, M., Hurvitz, A., Galeotti, M., Bacharach, E., Eldar, A., 2014. Identification of a novel RNA virus lethal to tilapia. Journal of Clinical Microbiology 52, 4137–4146. https://doi.org/10.1128/JCM.00827-14

FAO, 2022. The State of World Fisheries and Aquaculture 2022. Towards Blue Transformation. FAO. https://doi.org/10.4060/cc0461en

Feil, E.J., Li, B.C., Aanensen, D.M., Hanage, W.P., Spratt, B.G., 2004. eBURST: Inferring Patterns of Evolutionary Descent among Clusters of Related Bacterial Genotypes from Multilocus Sequence Typing Data. Journal of Bacteriology 186, 1518–1530. https://doi.org/10.1128/JB.186.5.1518-1530.2004

Feito, J., Araújo, C., Gómez-Sala, B., Contente, D., Campanero, C., Arbulu, S., Saralegui, C., Peña, N., Muñoz-Atienza, E., Borrero, J., del Campo, R., Hernández, P.E., Cintas, L.M., 2022. Antimicrobial activity, molecular typing and in vitro safety assessment of *Lactococcus garvieae* isolates from healthy cultured rainbow trout (*Oncorhynchus mykiss*, Walbaum) and rearing environment. LWT 162, 113496. https://doi.org/10.1016/j.lwt.2022.113496

Felsenstein, J., 1985. Confidence limits on phylogenies: An approach using the bootstrap. Evolution (N Y) 39, 783–791. https://doi.org/10.1111/j.1558-5646.1985.tb00420.x

Ferrario, C., Ricci, G., Milani, C., Lugli, G.A., Ventura, M., Eraclio, G., Borgo, F., Fortina, M.G., 2013. *Lactococcus garvieae*: Where is it from? A first approach to explore the evolutionary history of this emerging pathogen. PLoS ONE 8. https://doi.org/10.1371/journal.pone.0084796

Figueiredo, H.C.P., Nobrega Netto, L., Leal, C.A.G., Pereira, U.P., Mian, G.F., 2012. *Streptococcus iniae* outbreaks in Brazilian Nile tilapia (*Oreochromis niloticus* L:) farms. Brazilian Journal of Microbiology 43, 576–580. https://doi.org/10.1590/S1517-83822012000200019

Fox, J.G., Yan, L.L., Dewhirst, F.E., Paster, B.J., Shames, B., Murphy, J.C., Hayward, A., Belcher, J.C., Mendes, E.N., 1995. *Helicobacter bilis* sp. nov., a Novel *Helicobacter* Species Isolated from Bile, Livers, and Intestines of Aged, Inbred Mice, JOURNAL OF CLINICAL MICROBIOLOGY.

Fukushima, H.C.S., Leal, C.A.G., Cavalcante, R.B., Figueiredo, H.C.P., Arijo, S., Moriñigo, M.A., Ishikawa, M., Borra, R.C., Ranzani-Paiva, M.J.T., 2017. *Lactococcus garvieae* outbreaks in Brazilian farms Lactococcosis in *Pseudoplatystoma* sp. – development of an autogenous vaccine as a control strategy. Journal of Fish Diseases 40, 263–272. https://doi.org/10.1111/jfd.12509

Gibello, A., Galán-Sánchez, F., Blanco, M.M., Rodríguez-Iglesias, M., Domínguez, L., Fernández-Garayzábal, J.F., 2016. The zoonotic potential of *Lactococcus garvieae*: An overview on microbiology, epidemiology, virulence factors and relationship with its presence in foods. Research in Veterinary Science. https://doi.org/10.1016/j.rvsc.2016.09.010

Goodman, L.B., Lawton, M.R., Franklin-Guild, R.J., Anderson, R.R., Schaan, L., Thachil, A.J., Wiedmann, M., Miller, C.B., Alcaine, S.D., Kovac, J., 2017. *Lactococcus petauri* sp. nov., isolated from an abscess of a sugar glider. International Journal of Systematic and Evolutionary Microbiology 67, 4397–4404. https://doi.org/10.1099/ijsem.0.002303

Karami, E., Alishahi, M., Molayemraftar, T., Ghorbanpour, M., Tabandeh, M.R., Mohammadian, T., 2019. Study of pathogenicity and severity of *Lactococcus garvieae* isolated from rainbow trout (*Oncorhynchus mykiss*) farms in Kohkilooieh and Boyerahmad province. Fisheries and Aquatic Sciences 22. https://doi.org/10.1186/s41240-019-0135-2

Kawanishi, M., Yoshida, T., Yagashiro, S., Kijima, M., Yagyu, K., Nakai, T., Murakami, M., Morita, H., Suzuki, S., 2006. Differences between *Lactococcus garvieae* isolated from the genus *Seriola* in Japan and those isolated from other animals (trout, terrestrial animals from Europe) with regard to pathogenicity, phage susceptibility and genetic characterization. Journal of Applied Microbiology 101, 496–504. https://doi.org/10.1111/j.1365-2672.2006.02951.x

Kotzamanidis, C., Malousi, A., Bitchava, K., Vafeas, G., Chatzidimitriou, D., Skoura, L., Papadimitriou, E., Chatzopoulou, F., Zdragas, A., 2020. First Report of Isolation and Genome Sequence of *L. petauri* Strain from a Rainbow Trout Lactococcosis Outbreak. Current Microbiology 77, 1089–1096. https://doi.org/10.1007/s00284-020-01905-8

Kronvall, G., 2003. Determination of the real standard distribution of susceptible strains in zone histograms. International Journal of Antimicrobial Agents 22, 7–13. https://doi.org/10.1016/S0924-8579(03)00107-9

Kronvall, G., Smith, P., 2016. Normalized resistance interpretation, the NRI method: Review of NRI disc test applications and guide to calculations. APMIS. https://doi.org/10.1111/apm.12624

Leal, C.A.G., Tavares, G.C., Figueiredo, H.C.P., 2014. Outbreaks and genetic diversity of *Francisella noatunensis* subsp *orientalis* isolated from farm-raised Nile tilapia (*Oreochromis niloticus*) in Brazil. Genetics and Molecular Research 13, 5704–5712. https://doi.org/10.4238/2014.July.25.26

Lin, Y.S., Kweh, K.H., Koh, T.H., Lau, Q.C., Abdul Rahman, N.B., 2020. Genomic analysis of *Lactococcus garvieae* isolates. Pathology 52, 700–707. https://doi.org/10.1016/j.pathol.2020.06.009

Maiden, M.C.J., van Rensburg, M.J.J., Bray, J.E., Earle, S.G., Ford, S.A., Jolley, K.A., McCarthy, N.D., 2013. MLST revisited: The gene-by-gene approach to bacterial genomics. Nature Reviews Microbiology 11, 728–736. https://doi.org/10.1038/nrmicro3093

Martinovic, A., Cabal, A., Nisic, A., Sucher, J., Stöger, A., Allerberger, F., Ruppitsch, W., 2021. Genome Sequences of *Lactococcus garvieae* and Lactococcus petauri Strains Isolated from Traditional Montenegrin Brine Cheeses. Microbiology Resource Announcements 10. https://doi.org/10.1128/mra.00546-21

Miles, A.A., Misra, S.S., Irwin, J.O., 1938. The estimation of the bactericidal power of the blood. Journal of Hygiene 38, 732–749. https://doi.org/10.1017/S002217240001158X

Morick, D., Davidovich, N., Zemah-Shamir, Z., Bigal, E., Rokney, A., Ron, M., Blum, S.E., Fleker, M., Soto, E., Heckman, T.I., Lau, S.C.K., Wosnick, N., Tchernov, D., Scheinin, A.P., 2022. First Isolation and Characterization of *Streptococcus agalactiae* From a Stranded Wild Common Dolphin (*Delphinus delphis*). Front Mar Sci 9. https://doi.org/10.3389/fmars.2022.824071

Nishiki, I., Furukawa, M., Matui, S., Itami, T., Nakai, T., Yoshida, T., 2011. Epidemiological study on *Lactococcus garvieae* isolates from fish in Japan. Fisheries Science 77, 367–373. https://doi.org/10.1007/s12562-011-0332-0

Nishiki, I., Oinaka, D., Iwasaki, Y., Yasuike, M., Nakamura, Y., Yoshida, T., Fujiwara, A., Nagai, S., Katoh, M., Kobayashi, T., 2016. Complete genome sequence of nonagglutinating *Lactococcus garviea*e strain 122061 isolated from yellowtail in Japan. Genome Announcements 4. https://doi.org/10.1128/genomeA.00592-16

Ortega, C., Irgang, R., Valladares-Carranza, B., Collarte, C., Avendaño-Herrera, R., 2020. First identification and characterization of *Lactococcus garvieae* isolated from rainbow trout (*Oncorhynchus mykiss*) cultured in Mexico. Animals 10, 1–15. https://doi.org/10.3390/ani10091609

Osman, K.M., Al-Maary, K.S., Mubarak, A.S., Dawoud, T.M., Moussa, I.M.I., Ibrahim, M.D.S., Hessain, A.M., Orabi, A., Fawzy, N.M., 2017. Characterization and susceptibility of streptococci and enterococci isolated from Nile tilapia (*Oreochromis niloticus*) showing septicaemia in aquaculture and wild sites in Egypt. BMC Veterinary Research 13. https://doi.org/10.1186/s12917-017-1289-8

Ou, Y.J., Ren, Q.Q., Fang, S.T., Wu, J.G., Jiang, Y.X., Chen, Y.R., Zhong, Y., Wang, D.D., Zhang, G.X., 2020. Complete Genome Insights into *Lactococcus petauri* CF11 Isolated from a Healthy Human Gut Using Second- and Third-Generation Sequencing. Frontiers in Genetics 11. https://doi.org/10.3389/fgene.2020.00119

Pérez-Losada, M., Cabezas, P., Castro-Nallar, E., Crandall, K.A., 2013. Pathogen typing in the genomics era: MLST and the future of molecular epidemiology. Infection, Genetics and Evolution 16, 38–53. https://doi.org/10.1016/j.meegid.2013.01.009

Pot, B., Devriese, L.A., Ursi, D., Vandamme, P., Haesebrouck, F., Kersters, K., 1996. Phenotypic Identification and Differentiation of *Lactococcus* Strains Isolated from Animals. Systematic and Applied Microbiology 19, 213–222. https://doi.org/10.1016/S0723-2020(96)80047-6

Rao, S., Pham, T.H., Poudyal, S., Cheng, L., Nazareth, S.C., Wang, P., Chen, S., 2021. First report on genetic characterization, cell-surface properties and pathogenicity of *Lactococcus garvieae*, emerging pathogen isolated from cage-cultured cobia (*Rachycentron canadum*). Transboundary and Emerging Diseases. https://doi.org/10.1111/tbed.14083

Reed, L.J., Muench, H., 1938. Asimple method of estimating fifty per cent endpoints. American Journal of Epidemiology 27, 493–497. https://doi.org/10.1093/oxfordjournals.aje.a118408

Reguera-Brito, M., Galán-Sánchez, F., Blanco, M.M., Rodríguez-Iglesias, M., Domínguez, L., Fernández-Garayzábal, J.F., Gibello, A., 2016. Genetic analysis of human clinical isolates of *Lactococcus garvieae*: Relatedness with isolates from foods. Infection, Genetics and Evolution 37, 185–191. https://doi.org/10.1016/j.meegid.2015.11.017

Reichley, S.R., Ware, C., Steadman, J., Gaunt, P.S., García, J.C., LaFrentz, B.R., Thachil, A., Waldbieser, G.C., Stine, C.B., Buján, N., Arias, C.R., Loch, T., Welch, T.J., Cipriano, R.C., Greenway, T.E., Khoo, L.H., Wise, D.J., Lawrence, M.L., Griffin, M.J., Reichley, C.S., Brad Fenwick, E., 2017. Comparative Phenotypic and Genotypic Analysis of *Edwardsiella* Isolates from Different Hosts and Geographic Origins, with Emphasis on Isolates Formerly Classified as *E. tarda*, and Evaluation of Diagnostic Methods Downloaded from, Journal of Clinical Microbiology on October.

Russo, G., Iannetta, M., D’abramo, A., Mascellino, M.T., Pantosti, A., Erario, L., Tebano, G., Oliva, A., D’agostino, C., Trinchieri, V., Vullo, V., 2012. *Lactococcus garvieae* endocarditis in a patient with colonic diverticulosis: first case report in Italy and review of the literature. New Microbiologica 35, 495–501.

Saitou, N., Nei, M., 1987. The neighbor-joining method: a new method for reconstructing phylogenetic trees. Molecular Biology and Evolution 406–425. https://doi.org/10.1093/oxfordjournals.molbev.a040454

Shahin, K., Mukkatira, K., Yazdi, Z., Richey, C., Kwak, K., Heckman, T.I., Mohammed, H.H., Ortega, C., Avendaño-Herrera, R., Keleher, B., Hyatt, M.W., Drennan, J.D., Adkison, M., Griffin, M.J., Soto, E., 2022. Development of a quantitative polymerase chain reaction assay for detection of the aetiological agents of piscine lactococcosis. Journal of Fish Diseases. https://doi.org/10.1111/jfd.13610

Shahin, K., Veek, T., Heckman, T.I., Littman, E., Mukkatira, K., Adkison, M., Welch, T.J., Imai, D.M., Pastenkos, G., Camus, A., Soto, E., 2021. Isolation and characterization of *Lactococcus garvieae* from rainbow trout, *Onchorhyncus mykiss*, from California, USA. Transboundary and Emerging Diseases. https://doi.org/10.1111/tbed.14250

Subramaniam, K., Gotesman, M., Smith, C.E., Steckler, N.K., Kelley, K.L., Groff, J.M., Waltzek, T.B., 2016. Megalocytivirus infection in cultured Nile tilapia *Oreochromis niloticus*. Diseases of Aquatic Organisms 119, 253–258. https://doi.org/10.3354/dao02985

Tamura, K., Nei, M., Kumar, S., 2004. Prospects for inferring very large phylogenies by using the neighbor-joining method. Proceedings of the National Academy of Sciences 101, 11030–11035. https://doi.org/10.1073/pnas.0404206101

Tamura, K., Stecher, G., Kumar, S., 2021. MEGA11: Molecular Evolutionary Genetics Analysis Version 11. Molecular Biology and Evolution 38, 3022–3027. https://doi.org/10.1093/molbev/msab120

Tavares, G.C., de Queiroz, G.A., Assis, G.B.N., Leibowitz, M.P., Teixeira, J.P., Figueiredo, H.C.P., Leal, C.A.G., 2018. Disease outbreaks in farmed Amazon catfish (*Leiarius marmoratus* x *Pseudoplatystoma corruscans*) caused by *Streptococcus agalactiae*, *S. iniae*, and *S. dysgalactiae*. Aquaculture 495, 384–392. https://doi.org/10.1016/j.aquaculture.2018.06.027

Teixeira, L.M., Merquior, V.L.C., Vianni, M. da C.E., Carvalho, M. da G.S., Fracalanzza, S.E.L., Steigerwalt, A.G., Brenner, D.J., Facklam, R.R., 1996. Phenotypic and Genotypic Characterization of Atypical *Lactococcus garvieae* Strains Isolated from Water Buffalos with Subclinical Mastitis and Confirmation of *L. garvieae* as a Senior Subjective Synonym of *Enterococcus seriolicida*. International Journal of Systematic Bacteriology 46, 664–668. https://doi.org/10.1099/00207713-46-3-664

Tejedor, J.L., Vela, A.I., Gibello, A., Casamayor, A., Domínguez, L., Fernández-Garayzábal, J.F., 2011. A genetic comparison of pig, cow and trout isolates of *Lactococcus garvieae* by PFGE analysis. Letters in Applied Microbiology 53, 614–619. https://doi.org/10.1111/j.1472-765X.2011.03153.x

Thiry, D., Billen, F., Boyen, F., Duprez, J.N., Quenault, H., Touzain, F., Blanchard, Y., Clercx, C., Mainil, J.G., 2021. Genomic relatedness of a canine *Lactococcus garvieae* to human, animal and environmental isolates. Research in Veterinary Science 137, 170–173. https://doi.org/10.1016/j.rvsc.2021.04.032

Torres-Corral, Y., Santos, Y., 2021. Clonality of *Lactococcus garvieae* isolated from rainbow trout cultured in Spain: A molecular, immunological, and proteomic approach. Aquaculture 545. https://doi.org/10.1016/j.aquaculture.2021.737190

Tsai, M.A., Wang, P.C., Liaw, L.L., Yoshida, T., Chen, S.C., 2012. Comparison of genetic characteristics and pathogenicity of *Lactococcus garvieae* isolated from aquatic animals in Taiwan. Diseases of Aquatic Organisms. https://doi.org/10.3354/dao02516

Wang, C.Y.C., Shie, H.S., Chen, S.C., Huang, J.P., Hsieh, I.C., Wen, M.S., Lin, F.C., Wu, D., 2007. *Lactococcus garvieae* infections in humans: Possible association with aquaculture outbreaks. International Journal of Clinical Practice 61, 68–73. https://doi.org/10.1111/j.1742-1241.2006.00855.x

