## Supplemental Table 1 for "Emerging fish pathogens *Lactococcus petauri* and *L. garvieae* in Nile tilapia (*Oreochromis niloticus*) farmed in Brazil"

Supplementary material

Table S1. *Lactococcus petauri* and *L. garvieae gyrB* gene sequences used in the phylogenetic analysis

| Isolate ID | Species | Accession number |
|---|---|---|
| 48/20-02 | *L. petauri* | Submitted to NCBI |
| 48/20-05 | *L. petauri* | Submitted to NCBI |
| 48/20-08 | *L. petauri* | Submitted to NCBI |
| 48/20-09 | *L. petauri* | Submitted to NCBI |
| 48/20-10 | *L. petauri* | Submitted to NCBI |
| 48/20-11 | *L. petauri* | Submitted to NCBI |
| 48/20-12 | *L. petauri* | Submitted to NCBI |
| 48/20-14 | *L. petauri* | Submitted to NCBI |
| 48/20-16 | *L. petauri* | Submitted to NCBI |
| 30/19-01 | *L. garvieae* | Submitted to NCBI |
| 05/21-28 | *L. garvieae* | Submitted to NCBI |
| 67/21-01 | *L. petauri* | Submitted to NCBI |
| 67/21-05 | *L. petauri* | Submitted to NCBI |
| 68/21-01 | *L. petauri* | Submitted to NCBI |
| 68/21-05 | *L. petauri* | Submitted to NCBI |
| 69/21-03 | *L. petauri* | Submitted to NCBI |
| 69/21-07 | *L. petauri* | Submitted to NCBI |
| 67/21-15 | *L. petauri* | Submitted to NCBI |
| 67/21-18 | *L. petauri* | Submitted to NCBI |
| 03/22-02 | *L. petauri* | Submitted to NCBI |
| 03/22-09 | *L. petauri* | Submitted to NCBI |
| 04/22-01 | *L. petauri* | Submitted to NCBI |
| 04/22-07 | *L. petauri* | Submitted to NCBI |
| 33/22-01 | *L. petauri* | Submitted to NCBI |
| 33/22-03 | *L. petauri* | Submitted to NCBI |
| 34/22-05 | *L. petauri* | Submitted to NCBI |
| 34/22-06 | *L. petauri* | Submitted to NCBI |
| 46/22-01 | *L. petauri* | Submitted to NCBI |
| 46/22-05 | *L. petauri* | Submitted to NCBI |
| 62/22-02 | *L. petauri* | Submitted to NCBI |
| 62/22-04 | *L. petauri* | Submitted to NCBI |

| | | |
|---|---|---|
| 94/22-03 | *L. petauri* | Submitted to NCBI |
| B1726 | *L. petauri* | NZ_CP094882.1* |
| PAQ102015-99 | *L. petauri* | NZ_LXWL01000001.1* |
| CF11 | *L. petauri* | NZ_CP045924.1* |
| 159469 | *L. petauri* | NZ_MUIZ01000001.1* |
| ATCC49156 | *L. garvieae* | NC_015930.1* |
| Lg2 | *L. garvieae* | NC_017490.1* |
| JJJN1 | *L. garvieae* | NZ_CP026502.1* |

*Sequence from complete genome.
