## Supplemental Table 2 for "Emerging fish pathogens *Lactococcus petauri* and *L. garvieae* in Nile tilapia (*Oreochromis niloticus*) farmed in Brazil"

Supplementary material

Table S2. Primers used in the MLST analysis

| Gene /Product | Primer pairs (5′-3′) | Annealing temperature (°C) | Amplicon (bp) |
|---|---|---|---|
| *atp*A (α-subunit of ATP synthase) | F: TAYRTYGGKGAYGGDATYGC<br>R: CCRCGRTTHARYTTHGCYTG | 56 | 1180 |
| *tuf* (elongation factor EF-Tu) | F: ATATGCGGCCGCCATYGGHCACGTBGACCA<br>R: AAAATATGCGGCCGCTCNCCNGGCATNACCAT | 56 | 1080 |
| *als* (α-acetolactate synthase) | F: ATTCGGCTCAGACTTAGTTG<br>R: TTCAGCTGCTTCAACATCAA | 58 | 1076 |
| *gap*C (glyceraldehyde-3-phosphate dehydrogenase) | F: AAGTTGGTATTAACGGTTTCG<br>R: AAGTGTACGAACGAGGTTAG | 56 | 974 |
| *gal*P (galactose permease) | F: TGGGGAAAATTTAAACCTTGG<br>R: ATCATCAGAACGGCTGGAAG | 58 | 1070 |
| *gyr*B (DNA gyrase β-subunit) | F: CATGCTGGTGGTAAATTTGG [a]<br>R: GTCATCCATTTCTCCTAAACC | 58 | 1464 |
| *rpo*C (RNA polymerase β'-subunit) | F: TTGGTCCACAAAAGGACTGG [a]<br>R: TCACGTCCTTTTGCTTCCAT | 58 | 1377 |

**Reference**

Ferrario, C., Ricci, G., Milani, C., Lugli, G.A., Ventura, M., Eraclio, G., Borgo, F., Fortina, M.G., 2013. Lactococcus garvieae: Where is it from? A first approach to explore the evolutionary history of this emerging pathogen. PLoS ONE 8. https://doi.org/10.1371/journal.pone.0084796
